# Variability in white matter structure relates to hallucination proneness

**DOI:** 10.1101/2021.06.10.447901

**Authors:** Joseph F. Johnson, Michael Schwartze, Michel Belyk, Ana P. Pinheiro, Sonja A. Kotz

## Abstract

Hallucinations are a prominent transdiagnostic psychiatric symptom but are also prevalent in individuals who do not require clinical care. Moreover, persistent psychosis-like experience in otherwise healthy individuals may be related to increased risk to transition to a psychotic disorder. This suggests a common etiology across clinical and non-clinical individuals along a multidimensional psychosis continuum that may be detectable in structural variations of the brain. The current diffusion tensor imaging study assessed healthy individuals to identify possible differences in white matter associated with hallucination proneness (HP). This approach circumvents potential confounds related to medication, hospitalization, and disease progression common in clinical individuals. We determined how HP relates to white matter integrity in selected association, commissural, and projection fiber pathways putatively linked to psychosis. Increased HP was associated with enhanced fractional anisotropy (FA) in the right uncinate fasciculus, the right anterior and posterior arcuate fasciculus, and the corpus callosum. Although FA in cortico-cerebellar pathways revealed no relationship, streamline quantity between the left cerebellum and the right motor cortex positively correlated with HP. These findings support the notion of a psychosis continuum, providing first evidence of structural white matter variability associated with HP in healthy individuals. Furthermore, alterations in the targeted pathways likely indicate an association between HP-related structural variations and the putative salience and attention mechanisms that these pathways subserve.

## Introduction

Hallucinations are externally attributed percepts in the absence of corresponding sensory input^1^. They are common in multiple psychiatric and neurological conditions but also occur in the general population^2,3,4,5^. Consequently, hallucinations are defined as a contributing factor to an extended psychosis phenotype, expressed as a multidimensional continuum rather than categorical symptomatology^6,7,8^. Hallucinatory percepts occur in any sensory modality, yet auditory verbal hallucinations (AVH) are the most common with a lifetime prevalence of 6-13% in healthy individuals^9,10,11^ and have received substantial scientific interest. Although many “healthy hallucinators’’ remain diagnosis free, increased hallucination proneness (HP) increases the risk to transition to psychosis^12,13^. Shared environmental and familial risk factors in clinical and non-clinical individuals suggest that psychosis-like experiences, including hallucinations, are associated with a common etiology^6,7,8^. Although the transdiagnostic approach to psychotic experience is well established and a putative common etiology is proposed across the population, the majority of neural substrates of hallucination research has been limited to schizophrenia. Functional magnetic resonance imaging (fMRI) in this clinical group provides strong evidence for miscommunication within and between functional brain networks^14,15,16,17^. However, schizophrenia research lacks consensus on how the expression of an etiological continuum manifests in white matter pathways connecting dispersed brain regions. As multiple factors might contribute to incongruent findings in clinical groups (i.e., antipsychotic medication, duration of illness, and low sample sizes, inconsistent diagnostic categories), further insight into the etiology of psychosis in the general population is needed^18,19,20^. The current DTI study addressed this need and investigated if proneness to hallucinate is associated with detectable variability in the brain’s white matter connections of healthy individuals.

Here the white matter pathways were restricted to tracts commonly linked to a hallucinatory experience in patient groups. Importantly, each of these pathways plays a putative role in cognitive or neural mechanisms altered by hallucinations and are not limited to deficits specific to clinical profiles. The first target was the uncinate fasciculus (UF), a long-range association fiber pathway bridging the anterior temporal cortex and temporal poles with the orbitofrontal and inferior frontal cortex, and connecting the amygdaloid body to the hippocampal formation^21,22,23^ (Figure 1B). As this pathway links regions involved in memory retrieval and processing salient or emotional sensory stimuli^24,25,26^, it is particularly important in the operant conditioning for successful interaction with the external world^27^. Due to the broad applicability of UF function, it is not surprising that structural abnormalities in this pathway are reported in multiple psychiatric, neurological, and developmental conditions^25^. Within schizophrenia, alterations in the UF often reveal a reduction of diffusion directionality^28,29,30,31^, further exacerbated by the severity of negative symptoms^28,32,33,34^. Loss of UF integrity associated with negative symptom severity may therefore signify hyposalient processing of environmental cues. Conversely, although clinical subgroups with AVH also exhibit a decreased directionality of UF diffusion^35,36,37,38^, studies investigating positive symptom severity, including hallucinations, have sometimes reported a positive correlation^39,40,41^. Recent AVH theories have ascribed this variability in UF anisotropy to hypersalient processing of irrelevant stimuli or inner speech signals^17^. However, it remains unknown if the underlying UF white matter alterations, associated with this function, extend to hallucinations in any modality. We addressed this question by assessing the relationship between the HP spectrum in the general population and UF anisotropy.

**FIGURE 1.**
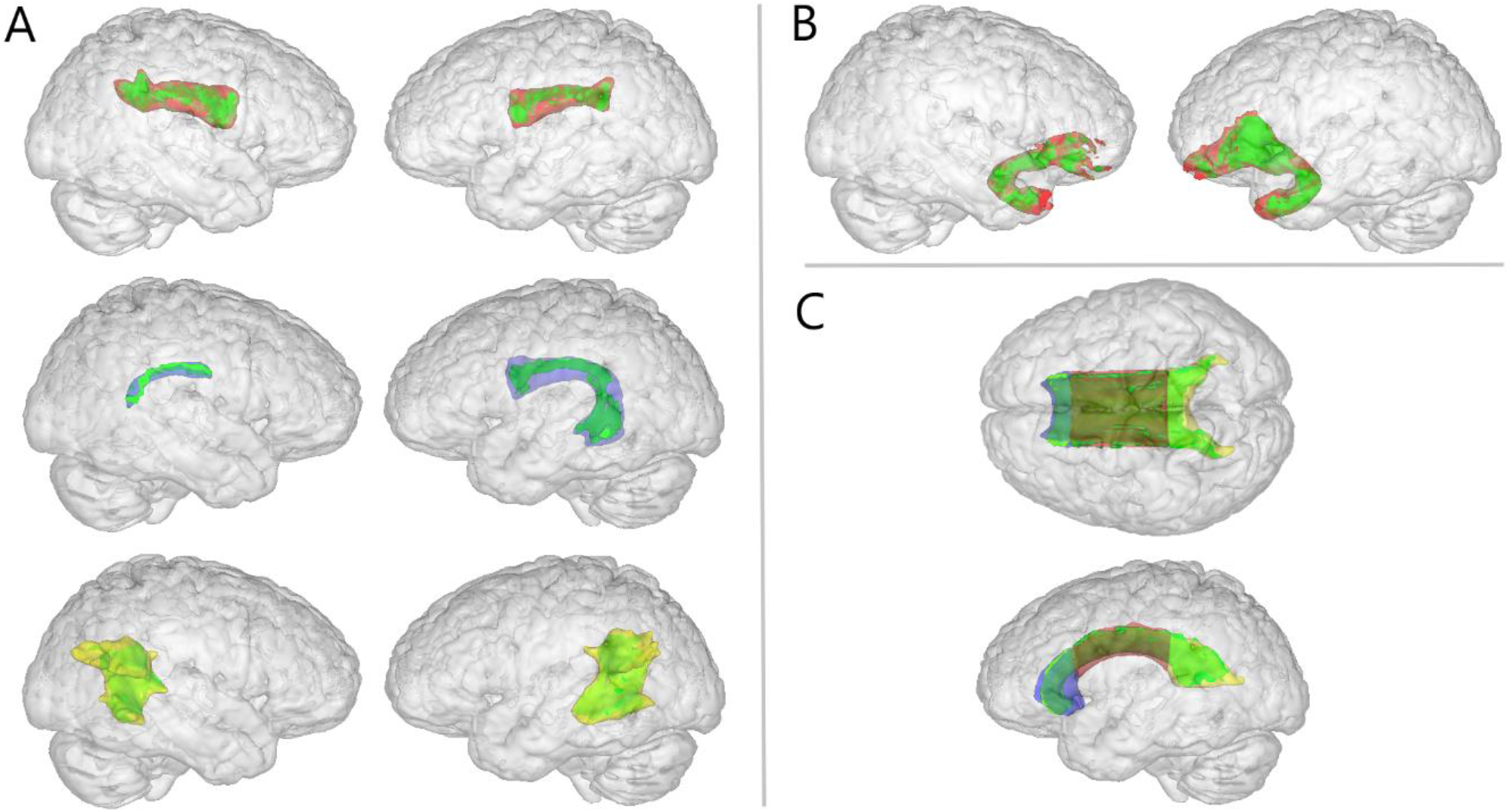
REGION OF INTEREST WHITE MATTER PATHWAYS: Green = voxels from group mean fractional anisotropy skeleton used in the correlation analysis; A. NATRAINLAB atlas perisylvian network (arcuate fasciculus) masks^52^: red = anterior segment, blue = longitudinal segment, yellow = posterior segment; B. red = JHU-white matter tractography atlas uncinate fasciculus masks; C. ICBM-DTI-81 white matter atlas corpus callosum: red = body, blue = genus, yellow = splenium.

The second target was the arcuate fasciculus (AF), a fronto-temporo-parietal (FT-P) association pathway of long and short fibers connecting regions implicated in speech planning and processing (Figure 1A). Due to its role in speech, the AF has been hypothesized to play a major role in auditory verbal hallucinations^15,42^. It has been posited that reduced communication between prefrontal speech planning regions and the auditory cortex during the monitoring of inner speech results in the perception of inner speech being attributed to an external source^43,44,45^. Although DTI evidence of reduced anisotropy in schizophrenia patients who hear voices supports this hypothesis^35,36,37,38,46,47,48,49,50^, there is also some evidence for an increase^35,51^. Furthermore, although a negative correlation between AVH severity and AF anisotropy has been reported^38^, both AVH^48,51^ and positive symptoms severity^51^ have also been positively correlated. Notably, in these studies there has been inconsistency regarding where in the AF alterations are observed, which may account for some variability in the respective findings. Fiber tracking has established a subdivision of the AF into segments that connect to different regions of the FT-P perisylvian pathway, allowing for a more fine-grained characterization of altered information transfer^52^. These segments include intra-hemispheric connections between the inferior frontal gyrus (IFG) and superior temporal gyrus (STG), the IFG and inferior parietal lobe (IPL), and the STG and IPL. As the IPL and temporal parietal junction connected to the AF have also been associated with a deficit in sensory self-other attribution in psychosis, this pathway may also be affected by hallucinations in other sensory modalities^53,54^. Accordingly, we segmented the AF to specify which aspects of FT-P processing may link to increased HP.

Third, we assessed a relationship of HP with the corpus callosum (CC), the largest collection of white matter fiber bundles in the brain (Figure 1C). Abnormal inter-hemispheric communication via the CC is often reported in schizophrenia and associated with a decrease in lateralized functions ascribed to a dominant hemisphere^17^. This is particularly relevant for AVH, as speech and language functions typically rely on asymmetrical processing in specialized functional regions of the auditory cortices^55^. The lateralization of these functions is affected by the degree of information transfer of the interhemispheric auditory pathway (IAP) joining left and right hemispheric homologues^56^. Previous DTI research in schizophrenia reported both FA increases^35,57,58^ and decreases^35,38,51,59,60,61^ associated with auditory hallucinations in CC structure. However, both AVH^58^ and positive symptom severity^51,62^ have shown a positive correlation with FA in the CC. Due to high variability in the specific localization of reported CC findings, we performed analyses of white matter anisotropy for individual segments^63^, including the genus, body, and splenium. This allowed clarifying if HP-related differences in CC white matter structure are specific to the IAP or associated with a general variability in interhemispheric communication.

Lastly, we investigated the relationship between HP and white matter pathways connecting the neocortex to the cerebellum. Unlike the functional aspects discussed for the neocortical pathways of interest, hallucinations associated with these tracts require the self-production of actual sensory feedback. Within this system, internal models of sensorimotor transformations are learned and updated via interaction with the environment^64^. Transformations of an action plan to expected sensory outcome (forward models) are compared to actual sensory feedback to both determine how our actions affect sensation, and how we must adapt our actions to unexpected sensory influences to reach a desired outcome. It is hypothesized that by this differentiation between the internal and external world, a sense of agency emerges^65^. This perspective suggests that as a motor command is sent to the periphery, the forward model associated with the outgoing command (efference copy) is sent in parallel to the cerebellum^66,67^. There, reafferent sensory feedback signals are continuously compared and discrepancies are communicated back via closed loops with the neocortex to inform about a need to update or adapt. Although the cerebellar system is implicated in hallucinatory experiences and the loss of distinction between sensation as internally or externally generated, it is still not known where exactly the breakdown in the system might occur. Erroneous transmission of the efference copy via the cortico-ponto-cerebellar (CPC) pathway may lead to sensory feedback being perceived as unexpected and external, whereas poor error signaling via the cerebello-thalamo-cortical (CTC) pathway may lead to reduced awareness of environmental influence on our feedback^68^. We therefore assessed white matter structure both within CPC and CTC pathways forming a reciprocal loop with the primary motor cortex (M1).

Using the Launay Slade Hallucination Scale (LSHS) as a reliable self-report measure of HP^69^, we examined how proneness in the general population might link to variations in white matter structure in the selected white matter tracts of interest. For the sake of comparison with existing and future research, we employed region of interest (ROI) analyses of mean FA within well-established atlas-based masks of association and commissural fiber pathways. Furthermore, due to interindividual variability in specific localization of reciprocal cortico-cerebellar-cortical loops, we employed probabilistic tractography to model the CPC and CTC projections to investigate possible variations in either efference copy or error signaling pathways. Although schizophrenia research has noted a general decrease in the directionality of diffusion in white matter pathways, reports of positive correlation in positive symptom and hallucination severity are increasingly prominent. Therefore, we hypothesized these pathways should reveal greater anisotropy as HP increases, accounting for aberrant communication between proximal regions. In doing so, we present findings that characterize structural disparities associated with the etiological continuum that may be linked to a heightened risk to transition to psychotic disorder. Furthermore, we aimed to bridge the gap between HP-related white matter variability and theoretical underpinnings of this complex phenomenology.

## Methods

### Participants

Fifty-two participants were recruited through the SONA system and social media channels at Maastricht University, the Netherlands. All had normal (or corrected-to-normal) hearing and vision. Participants provided informed consent and received university study credit for compensation. Exclusion criteria were any history of psychotic disorder, neurological impairment, metal implants, previous traumatic brain injury, claustrophobia, or pregnancy. Robust statistics determined the age of one participant as an outlier, leading to the removal of this dataset from further analyses. One additional participant was excluded due to a scanning artifact. Of the remaining 50 participants (35 female), the average age was 22.52 years (SD 4.27; range 18 to 34). The study was approved by the Ethical Review Committee of the Faculty of Psychology and Neuroscience at Maastricht University (ERCPN-176_08_02_2017).

### Hallucination proneness

To measure HP, all participants filled in the revised-LSHS^69^. This five-point Likert scale self-report questionnaire consists of 16 questions, with items targeting levels of tactile, sleep-related, visual, and auditory hallucinations as well as vivid thoughts and daydreaming. To test for covariation of these data with measures of white matter structure, we calculated total LSHS scores, comprising all 16 items for each participant as a measure of overall HP. Furthermore, we conducted an exploratory analysis on a subset of 3 auditory items of the LSHS to determine if results of white matter correlation were driven specifically by an auditory-related psychosis-like experience. This subset was previously validated as loading under a single factor through principal component analyses^69^.

### Data acquisition

Neuroimaging data were collected using a Siemens 3T Magnetom Prisma Fit MRI scanner equipped with a 32-channel head coil (Siemens Healthcare, Erlangen, Germany) at the Scannexus facilities (Maastricht, the Netherlands). For each participant, a T1-weighted single-shot echoplanar imaging (EPI) sequence was collected using a repetition time (TR) of 2250 ms, 2.21 ms echo-time (TE), 256 mm field of view (FoV), 192 slices interleaved, 1.0 mm slice thickness (voxel size 1.0 mm3), and anterior-posterior phase encoding direction. In the same session, diffusion weighted images were recorded using a 8400 ms TR, 53 ms TE, 204 mm FoV, 87 slices interleaved, 1.5 mm slice thickness (voxel size 1.5 mm3), parallel imaging (GRAPPA) with factor 2, 30 diffusion-encoding gradients with b-value 1000 s/mm2, one b-value 0 (no diffusion weighting), and anterior-posterior phase encoding direction. The sequence was then repeated in the reverse phase encoding direction to correct for susceptibility-induced distortion. Total acquisition time was about 13 minutes.

### Data pre-processing

Imaging data were converted from DICOM to 4D NIFTI using the MRIcron software Dcm2Nii conversion tool (https://www.nitrc.org/projects/mricron/). Files containing b-value and b-vector data were retrieved simultaneously by Dcm2Nii file conversion. DTI data (pre-)processing was performed using the FMRIB Diffusion Toolbox (FDT) of FSL version 6.0.3 software (FMRIB Software Library, Oxford, United Kingdom, http://www.fmrib.ox.ac.jk/fsl)^70^. Topup was used to estimate the susceptibility-induced off-resonance field for both anterior-posterior and reverse phase-encode blips and to form a single corrected image^71^, non-brain tissue was then removed using the brain extraction tool (BET)^72^, and finally eddy current-induced distortions and participant movements were corrected^73^.

### Tractography

Due to high intersubject variability in the trajectory of connections between the cortex and cerebellum, tractography was used to model four projection fiber pathways for subsequent extraction of mean FA, representing left and right CPC and CTC. Crossing fibers analysis was conducted for each participant in their own subjective diffusion space using the pre-processed diffusion weighted and brain-extracted mask images. Using BEDPOSTX from the FSL FDT^74^, the number of crossing fibers within each voxel of the brain was determined and modeled, using default settings (number of fibers = 2, weight = 1, burn in = 1000).

For the CPC pathways, a seed mask was set in the left or right primary M1 with waypoints in the cerebral peduncle, pons, middle cerebellar peduncle and terminating in the contralateral cerebellar cortex. Using the FSL FLIRT toolbox^75^, all seed and waypoints were determined using standard atlas-based masks and registered to the individual structural space of each dataset (Sup. Table 1B). An exclusion mask for each participant restricted any erroneous tracking through the midline prior to decussation in the pons, as well as fibers passing through the ipsilateral cerebellum. Importantly, tracking through the cerebral peduncle was restricted to include only streamlines in the posterior corticopontine segmentation of the crus cerebri. As many fibers funnel and cross through this narrow structure, this restriction was essential to avoid both the tracking of parallel corticospinal fibers coming from the same region of M1 and corticopontine fibers from the prefrontal cortex. The CTC pathways were seeded in the entire left or right cerebellar cortex with waypoints in the ipsilateral superior cerebellar peduncle, the contralateral ventrolateral thalamic nuclei, and terminating in the contralateral M1. Exclusion masks for each participant restricted erroneous tracking through the midline or the middle cerebellar peduncle. Finally, a transformation matrix was created for each dataset to convert all masks in structural space to diffusion space for tractography using linear registration in FLIRT.

Tractography was completed for each participant with the pre-processed diffusion weighted images and BEDPOSTX output using PROBTRACKX_gpu^76^. Default settings were used (5000 samples, 2000 steps, 0.5 step-length, 0.02 curvature threshold, 0.01 fiber threshold, 0.0 distance threshold). A loop check function was applied to avoid including streamlines that loop back onto the pathway of interest. The modeled pathways were then converted into the structural space of each participant and binarized to create masks for subsequent FA extraction. In addition, for each pathway the number of streamlines was extracted as a measure of connectivity from seed to end mask. This connectivity measure was then related to the LSHS HP and the auditory items subset using Pearson’s correlation.

### Correlation analyses

The diffusion tensor was fitted to the images with DTIFIT, while diffusion tensor parameters were estimated for each voxel via extracted eigenvectors and eigenvalues in three directions (λ1, λ2, λ3). Measures characterizing diffusivity such as FA and mean diffusivity (MD) were produced automatically via DTIFIT, while axial diffusivity (AD: λ║ ≡ λ1 > λ2, λ3) and radial diffusivity (RD: λ┴ ≡ (λ2 + λ3)/2) were calculated manually. FA is a measure of the proportional magnitude of directional movement of water along axonal fibers, which is commonly referred to as an indicator of white matter integrity^77^. It may be enhanced by various factors such as increases of parallel diffusion, restriction of perpendicular diffusion, or a combination of these aspects. Therefore, in areas of significant FA differences, measures of MD, AD, and RD representing the magnitude of water diffusion over all directions, parallel, and perpendicular to the tract can help inform the neurobiological interpretation of FA^77^. Finally, the tract-based spatial statistics (TBSS) toolbox was used to erode remaining non-brain tissue, register, and warp all the images of all participants to a common space, and produce a mean FA skeleton^77^. This process was repeated with the non-FA TBSS function to produce whole-brain maps of mean MD, AD, and RD.

In addition to the cortico-cerebellar-cortical masks modeled via tractography, atlas-based white ROI were modeled in MNI standard space: left and right UF, left and right anterior, longitudinal, and posterior AF, and body, genu, and splenium of the CC (Sup. Table 1A). ROI mask-building was accomplished by multiplying the mean FA skeleton mask with atlas-based white matter tract segmentations. The probabilistic atlas-based selections were binarized to form mask images and were re-sampled to match FMRIB58 FA 1mm space. Finally, using individual FA maps the mean FA within each of the four projection fiber pathways connecting neocortical and cerebellar regions, as well as 8 association and 3 commissural fiber pathways connecting neocortical regions were extracted for each participant.

Bivariate Pearson correlation analyses were then conducted for each ROI, relating individual mean FA to the LSHS measure of overall HP. This process was repeated for the LSHS auditory-only subset of items. All correlation analyses were conducted in SPSS (Version 26). To account for multiple comparisons, a false discovery rate (FDR) Benjamini-Hochberg correction was applied with an adjusted alpha of 0.05. For ROIs reporting positive FA findings, further significant correlations between HP and MD, AD, and RD were reported.

## Results

### Hallucination Proneness

The mean HP score for the 50 participants was 19.86 (*SD* = 10.72; range 2 to 42) out of a possible 64 points. Using a Shapiro-Wilk test, the distribution of HP scores in the sample was not significantly different from the normal distribution (p = 0.078). However, as the test of normality neared rejection, a bootstrapping procedure (1000 samples) and confidence interval correction was adopted for all correlation analyses between white matter measures and HP. Pearson’s correlation analysis reported a strong positive correlation between the auditory item subscore and a subscore of all non-auditory items (*r*_(49)_ = 0.620), p < 0.001).

### Atlas-based Correlation Analyses

Significant positive correlations with LSHS hallucination proneness scores were found in the right anterior and posterior portions of the AF (Table 1A, Figure 2A), as well as in the right UF (Table 1B, Figure 2B). Likewise, all subsections of the CC (genu, body, and splenium) displayed a positive correlation with hallucination proneness scores (Table 1C, Figure 2C). For regions reporting significant correlation to FA, the right posterior AF and splenium of the CC also produced a positive correlation with AD (*r*_(49)_ = 0.401, FDR-*p* = 0.012; *r*_(49)_ = 0.378, FDR-*p* = 0.020). The genus of the CC also correlated negatively with MD (*r*_(49)_ = −0.337, *FDR-p* = 0.036) and RD (*r*_(49)_ = −0.319, FDR-*p* = 0.036). No atlas-based pathways reported a correlation between mean FA and the LSHS auditory item subset (Sup. Table 2A).

**TABLE 1.** ATLAS-BASED ROI CORRELATION RESULTS: Pearson’s correlation analysis results between mean fractional anisotropy and LSHS composite score of hallucination proneness. Benjamini-Hochberg FDR-*p* correction for multiple comparisons (* = *p* < 0.05 threshold for significance). Bias-corrected confidence intervals (95%, bootstrapping with 1000 samples).

| Tract |  | *r* | 95% CI (bias-corrected) | FDR-*p* |
| --- | --- | --- | --- | --- |
| **A. Arcuate Fasciculus** |  |  |  |  |
| **Left** | **Longitudinal** | 0.218 | 0.005-0.435 | 0.156 |
|  | **Anterior** | 0.186 | -0.064-0.425 | 0.196 |
|  | **Posterior** | 0.261 | -0.007-0.490 | 0.092 |
| **Right** | **Longitudinal** | 0.197 | -0.045-0.428 | 0.187 |
|  | **Anterior** | 0.327 | 0.051-0.567 | 0.042* |
|  | **Posterior** | 0.401 | 0.119-0.672 | 0.021* |
| **B. Uncinate Fasciculus** |  |  |  |  |
| **Left** |  | 0.297 | 0.044-0.523 | 0.057 |
| **Right** |  | 0.362 | 0.026-0.665 | 0.036* |
| **C. Corpus Callosum** |  |  |  |  |
| **Genus** |  | 0.457 | 0.139-0.704 | 0.009* |
| **Body** |  | 0.331 | 0.079-0.586 | 0.042* |
| **Splenium** |  | 0.321 | 0.022-0.582 | 0.042* |

**TABLE 2.** TRACTOGRAPHY-BASED CORRELATION RESULTS: Pearson’s correlation analysis results between mean fractional anisotropy and LSHS composite score of hallucination proneness, and number of streamlines and LSHS composite score of hallucination proneness. Benjamini-Hochberg FDR-*p* correction for multiple comparisons (* = *p* < 0.05 threshold for significance). Bias-corrected confidence intervals (95%, bootstrapping with 1000 samples).

| Tract |  | *r* | 95% CI (bias-corrected) | FDR-*p* |
| --- | --- | --- | --- | --- |
| **Cortico-ponto-cerebellar pathway** |  |  |  |  |
| FA | Right M1 → Left CE | 0.162 | -0.099-0.400 | 0.573 |
|  | Left M1 → Right CE | 0.128 | -0.121-0.362 | 0.573 |
| # Streamlines | Right M1 → Left CE | -0.026 | -0.340-0.280 | 0.865 |
|  | Left M1 → Left CE | -0.129 | -0.363-0.137 | 0.524 |
| **Cerebello-thalamo-cortical pathway** |  |  |  |  |
| FA | Right CE → Left M1 | 0.093 | -0.189-0.358 | 0.573 |
|  | Left CE → Right M1 | 0.188 | -0.153-0.616 | 0.573 |
| # Streamlines | Right CE → Left M1 | 0.270 | -0.050-0.575 | 0.194 |
|  | Left CE → Right M1 | 0.420 | 0.153-0.616 | 0.028* |

**FIGURE 2.**
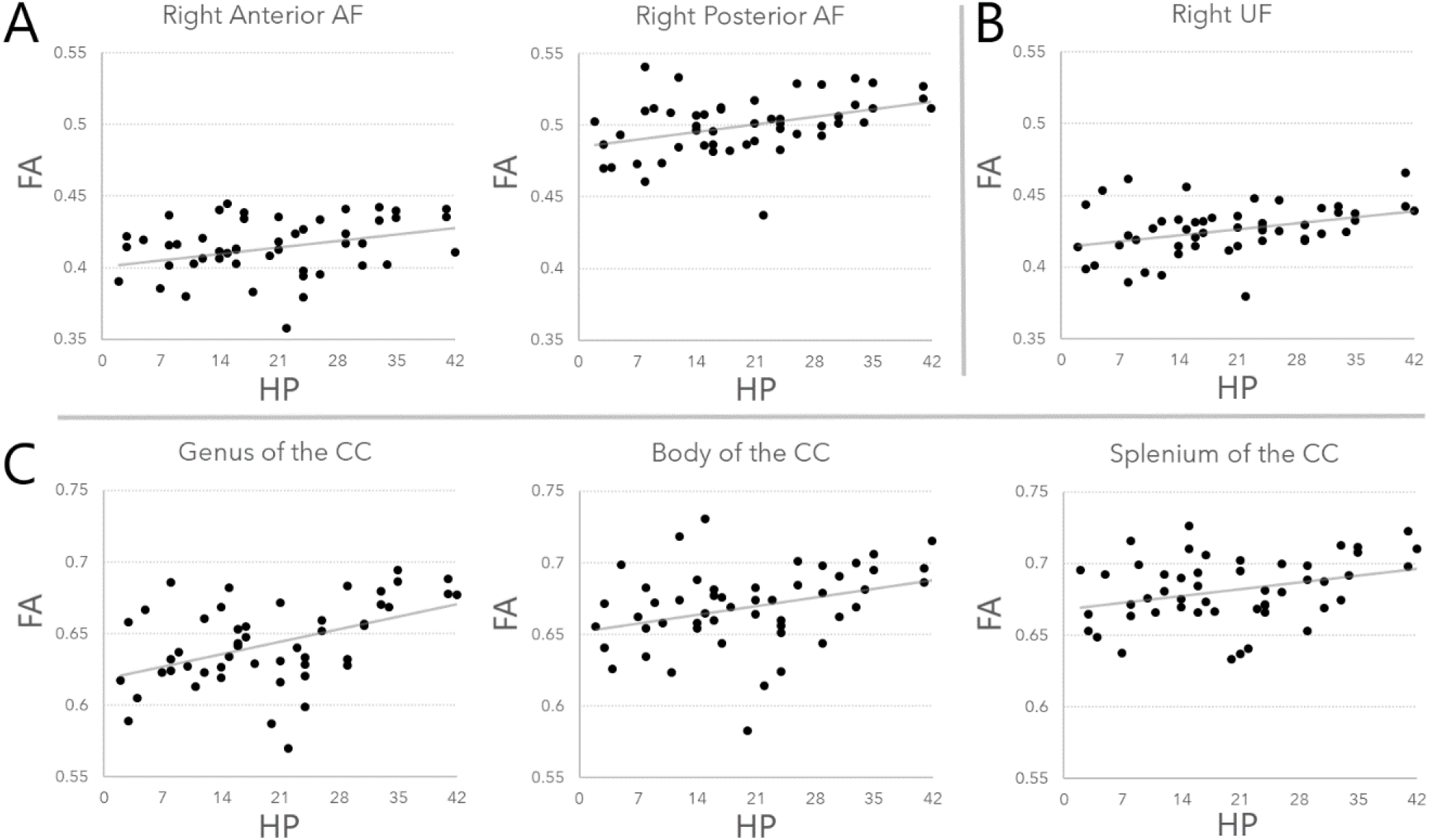
REGION OF INTEREST CORRELATION RESULTS: Pearson’s correlation between mean fractional anisotropy and LSHS composite score of hallucination proneness. A. Arcuate fasciculus: right anterior (*r*_(49)_ = 0.327, FDR-*p* = 0.042) and right posterior segments (*r*_(49)_ = 0.401, FDR-*p* = 0.021); B. Right Uncinate fasciculus (*r*_(49)_ = 0.362, FDR-*p* = 0.036); C. Corpus callosum: genus (*r*_(49)_ = 0.457, FDR-*p* = 0.009), body (*r*_(49)_ = 0.331, FDR-*p* = 0.042), splenium (*r*_(49)_ = 0.321, FDR-*p* = 0.042).

### Tractography-based Correlation Analyses

Successful tracking was completed in contralateral CPC pathways for 46 participants (Figure 3A). In the CTC pathways, the right cerebellum to the left M1 was successfully tracked for 39 participants, and the left cerebellum and right M1 for 40 (Figure 3B). For participants with successful tracking in both CTC, paired samples t-tests between number of streamlines of the left cerebellum to right M1 (*M* = 13.440, *SD* = 14.743) and right cerebellum to left M1 (*M* = 10.620, *SD* = 11.835) showed no significant difference (*t*_(33)_ = 1.083, FDR-*p* = 0.287). However, for the CPC there was a significant difference (*t*_(45)_ = −7.117, FDR-*p* < 0.001) with more streamlines between the right M1 and left cerebellum (*M* = 1247.478, *SD* = 1061.944) compared to the left M1 to right cerebellum (*M* = 190.700, *SD* = 183.833). None of the modeled cerebellar pathways revealed a significant correlation between HP and the extracted mean FA. However, the number of streamlines of only the CTC pathway displayed a significant positive correlation with HP, from the left cerebellar cortex to the right M1 (Figure 3C). No significant results were obtained for measures of white matter structure and the LSHS auditory item subset in any either CPC or CTC pathways (Sup. Table 2B).

**FIGURE 3.**
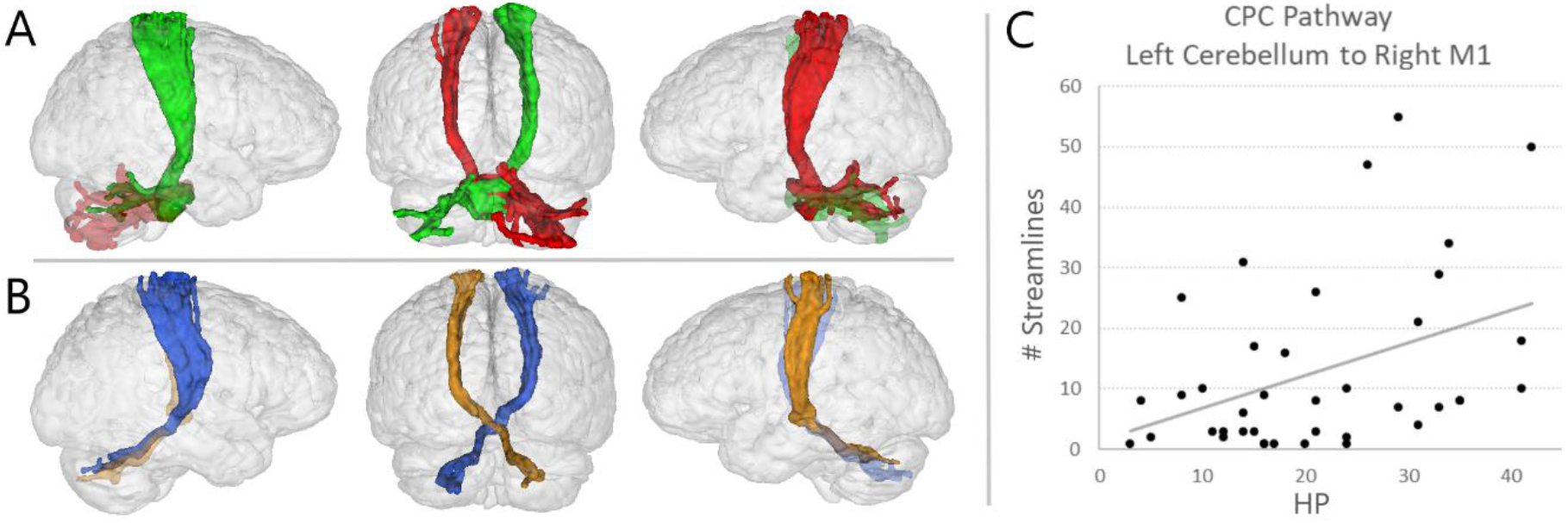
CORTICO-CEREBELLO-CORTICAL TRACTOGRAPHY AND CORRELATION RESULTS: A. Cortico-ponto-cerebellar pathway sample tracking (participant P08): green = right primary motor cortex to left cerebellum (221 streamlines), red = left primary motor cortex to right cerebellum (104 streamlines); B. Cerebello-thalamo-cortical pathway sample tracking (participant P08): blue = left cerebellum to right primary motor cortex (50 streamlines), orange = right cerebellum to left primary motor cortex (22 streamlines); C. Significant results from Pearson’s correlation analysis results between number of streamlines and LSHS composite score of hallucination proneness: cortico-ponto-cerebellar pathway from left cerebellum to right primary motor cortex (*r*_(39)_ = 0.420, FDR-*p* = 0.028).

## Discussion

The experience of hallucinations by clinical and non-clinical individuals suggests a common etiology along a psychosis continuum^6,7,8^. The current findings confirm that hallucination proneness covaries with structural white matter variability in healthy individuals. An increase in the directionality of diffusion was evident in right hemisphere fronto-temporal association and commissural pathways and in increased number of connections between the left cerebellum and the right motor cortex. These findings suggest that white matter pathways associated with psychosis putatively involved in salience, auditory, and sensory feedback networks are already affected in individuals who are prone to hallucinate but do not require clinical care.

### Memory/limbic networks and hypersalience

Although DTI research has indicated the involvement of UF in psychosis and psychosis-like experiences, there is no consensus regarding its relationship to phenomenology. While negative symptom severity has consistently shown a relationship with decreases in the directionality of UF diffusion^28,32,33,34^, positive symptoms have conversely provided some evidence for an increase^39,40,41^. Not only has this pathway shown variability within schizophrenia, but it has been linked to many cognitive functions and been implicated in several psychiatric, neurological, and developmental disorders^25,27^. Functionally divergent subdivisions of the UF may contribute to this disparity. These subdivisions may be differentially affected, leading to heterogeneous symptoms, or may be a common contributor to transdiagnostic symptoms^79,80^. Therefore, studies conducted in non-clinical groups may provide clearer findings, for example, regarding psychosis-like experiences in the general population. A large sample population-based study of non-clinical adolescents with psychotic symptoms confirmed patterns of increased directionality of diffusion in the UF^81^. Conversely, a decrease in UF white matter integrity was linked to the progression of psychosis pathology^82^. This indicates divergent contributions of UF structure integrity to symptom severity and disease progression. In line with non-clinical symptom-related findings, the current study reported FA of the right UF to be positively correlated with HP.

The UF carries internal connections of a memory and limbic system and plays an important role in the processing of salient environmental cues^21,22,23^. According to the triple-network model^83^, erroneous engagement of the salience network can disengage the default mode network and trigger active sensory processing via the central executive network^84^. For example, hypersalient processing of inner speech in the default mode can result in activation of speech processing regions and the perception of external speech in the form of AVH^15,16,38,85^. This disengagement of functional brain networks at rest is suggested more broadly to be associated with the misattribution of internally/externally generated stimuli^86^. Importantly, the network switching hypothesis has been related to an increased risk for psychosis or subthreshold positive symptoms including hallucinations^87,88^. The reported spectrum of increased white matter integrity in the healthy participants might therefore not only shed light on the general role of the UF in salience processing, but also provide a structural region of interest for future research on the risk for developing psychosis.

### Fronto-temporo-parietal networks top-down/bottom-up signals

The AF is commonly linked to AVH based on its putative role in self-monitoring of inner speech, during which predictions about sensory consequences from the IFG are sent to the auditory cortex, leading to the suppression of cortical responses to self-generated signals^15,42,43,44,45^. However, reports of increased^35,51^ and decreased^35,36,37,38,46,47,48,49,50,51^ FA in patients with AVH and variation in the precise location of AF differences necessitate further differentiation. Separate functions of three overlapping AF subdivisions could explain these conflicting findings. A direct long segment connects inferior frontal to posterior temporal, an anterior indirect segment connects inferior parietal to inferior frontal, and a posterior indirect segment connects posterior temporal to inferior parietal regions^52^. Some DTI studies have reported both increases and decreases in FA in separate segments of the FT pathway^35,51,89^. Furthermore, although the directionality of diffusion in schizophrenia patients with AVH provided mixed results, symptom severity of auditory hallucinations positively correlated with anisotropy^48,51,89^. The current findings of increased FA in the AF are in line with such links to symptom severity. However, as these variations were only found in the anterior and posterior segments of the right hemisphere, it is unlikely that the findings indicate an involvement of speech processing that has been ascribed to connections between the left IFG and STG.

Models of predictive inference^90^ alternatively suggest a more basic role of FT-P top-down predictive and bottom-up sensory input imbalance in a hallucinatory experience^91^. Due to a lack of inhibition, an influx of excitatory top-down inputs to sensory cortices results in overactivation in conditions of minimal bottom-up stimulation^92^. This leads to the development of false inference misattributions of internally generated predictions of inner speech that are perceived as real external voices. Therefore, although the weak prediction signaling of self-monitoring and strong priors of predictive inference accounts differ in the proposed mechanisms, both attribute hallucinations to an overactivation of sensory cortical regions^93^. This interpretation of increased top-down excitatory signals is in line with the current UF and AF HP-related increases in FA. Within the triple-network model, information transfer from inferior frontal hypersalient and central executive engagement resulting in activation of the temporal speech regions may correspond structurally with the anterior AF segment. Indeed, the right IFG has shown increased activity in monitoring unexpected features of auditory and voice feedback^94,95^. Further, the HP-related increases in the FA of the right posterior AF reported here suggest a contribution of structural connections to a sense of agency over external stimuli. The temporal parietal junction, a terminus point of the posterior AF, has been associated with variability in awareness and self-other attribution for neuropsychiatric symptoms such as hallucinations^53,54^. One limitation of the current results is that the reported atlas-based parcellations of the AF contain some overlap between the identified subregions. Therefore, although the long AF subsegment did not show a significant difference, we cannot rule out that some long association fibers joining IFG to STG are included across anterior and posterior AF ROIs.

### Interhemispheric miscommunication and hemispheric specialization

Contrary to theories of IAP involvement, the current study and previous AVH investigations show that variability in the anisotropy of CC commissural pathways is found across different subdivisions. Our findings indicate that the genu, body, and splenium are susceptible to greater anisotropy with increasing HP. Therefore, we suggest that the overall CC involvement is not solely resulting from increased communication of bilateral auditory regions associated with auditory hallucinations. Two opposing theories have been proposed to understand how CC structure may affect hemispheric specialization in regions affected by AVH^96^. In line with our findings of increased right AF integrity in the typically left-specialized language network, the excitatory model postulates that increased interhemispheric information transfer decreases hemispheric specialization. The inhibitory model suggests that increased interhemispheric communication maintains independent processing. An extension of the IMT proposed that top-down prefrontal cognitive control is preserved in healthy hallucinators^91^. This allows for the perceptual experience of hallucination without the belief that the voice comes from an external source. Therefore, this account is incongruent with our proposals of a role of top-down cognitive control associated with increasing HP. However, the observed increased genus of the CC anisotropy between frontal regions may support a compensatory role of right-specialized prefrontal cognitive control in healthy HP to counteract decreased specialization of the left-specialized language network.

### Cortico-cerebello-cortical loops and sensorimotor feedback error

The cerebellar role in sensorimotor feedback processing provides another key element to explain the phenomenology of hallucinatory experience^66,67^. As in models of perceptual inference, the proposed cerebellar mechanisms rely on sensory prediction. However, this system is evolved to facilitate successful interaction with the environment, and therefore requires actual sensory feedback for learning and updating forward models of expected sensory outcomes of action^97^. Cortico-cerebello-cortical circuits form closed-loops, where efference copies are propagated to the cerebellum via the pontine nuclei, compared to actual reafferent feedback, and resulting discrepancies are then signaled back to the cortex via the thalamus^98^. Incorrect functioning of this system can result in the loss of distinction between internally and externally generated sensation, however very few studies have attempted to link hallucination or the severity of positive psychosis symptoms to the structure of cerebellar pathways^39,99,100^. Moreover, these studies have been conducted exclusively in patient samples and employed varied methodologies that produced inconsistent results. Using tractography, we modelled and analyzed the cortico-cerebello-cortical loop between motor cortex and cerebellar cortex of each participant (Figure 1D). Results indicated no correlation of FA to measures of proneness to hallucinate. However, an increase in the number of streamlines correlated positively with HP in the cerebello-thalamo-cortical pathway, a connection with a putative involvement in prediction error signaling and/or updating of the forward model^68^. In the cortico-ponto-cerebellar pathway, no correlation with number of streamlines was found, suggesting no association to the propagation of an efference copy to the cerebellum.

### Conclusions

Hallucinations are a transdiagnostic phenomenon in multiple disorders and the general population. The results of the current study support an etiological continuum model of psychosis, as white matter pathways involved in salience (UF), perceptual inference (AF), and interhemispheric communication (CC) increase in the directionality of diffusion as the proneness to hallucinate rises in healthy individuals. All significant FA differences were found along the midline in the corpus callosum or limited to the right hemisphere, further suggesting variation in the lateralization of pathways in those more likely to hallucinate. Despite careful modelling of difficult to study cortico-cerebellar-cortical sensorimotor feedback loops of the human brain, FA in these pathways did not correlate with HP. However, a correlation of HP to the number of streamlines in the CTC tract implicated structural pathways involved in error-signaling and updating forward models. As hallucinatory experience is seen in different patient groups and the general population, we attempted to outline the putative roles of the affected white matter pathways in generalized brain mechanisms, as opposed to deficits reported in clinical groups. The current evidence suggests that variability in white matter structure associated with proneness to hallucinate may be related to mechanisms responsible for the attribution of salience and attention to sensory inputs, and not specific to language networks. Our findings support the contribution of a common etiology for HP across a continuum. We suggest that further research should reveal how this variability in structure may indicate a potential risk to transition to illness.

## References

1. Bentall, R. P. (1990). The illusion of reality: a review and integration of psychological research on hallucinations. Psychological bulletin, 107(1), 82.

2. Van Os, J., Hanssen, M., Bijl, R. V., & Ravelli, A. (2000). Strauss (1969) revisited: a psychosis continuum in the general population? Schizophrenia research, 45(1-2), 11–20.

3. Waters, F., & Fernyhough, C. (2017). Hallucinations: a systematic review of points of similarity and difference across diagnostic classes. Schizophrenia bulletin, 43(1), 32–43.

4. Rollins, C. P., Garrison, J. R., Simons, J. S., Rowe, J. B., O’Callaghan, C., Murray, G. K., & Suckling, J. (2019). Meta-analytic evidence for the plurality of mechanisms in transdiagnostic structural MRI studies of hallucination status. EClinicalMedicine, 8, 57–71.

5. Zhuo, C., Jiang, D., Liu, C., Lin, X., Li, J., Chen, G., … & Zhu, J. (2019). Understanding auditory verbal hallucinations in healthy individuals and individuals with psychiatric disorders. Psychiatry research, 274, 213–219.

6. Johns, L. C., & Van Os, J. (2001). The continuity of psychotic experiences in the general population. Clinical psychology review, 21(8), 1125–1141.

7. Johns, L. C. (2005). Hallucinations in the general population. Current psychiatry reports, 7(3), 162–167.

8. Myin-Germeys, I., Krabbendam, L., & van Os, J. (2003). Continuity of psychotic symptoms in the community. Current Opinion in Psychiatry, 16(4), 443–449.

9. Beavan, V., Read, J., & Cartwright, C. (2011). The prevalence of voice-hearers in the general population: a literature review. Journal of Mental Health, 20(3), 281–292.

10. Linscott, R. J., & Van Os, J. (2013). An updated and conservative systematic review and meta-analysis of epidemiological evidence on psychotic experiences in children and adults: on the pathway from proneness to persistence to dimensional expression across mental disorders. Psychological medicine, 43(6), 1133.

11. McGrath, J. J., Saha, S., Al-Hamzawi, A., Alonso, J., Bromet, E. J., Bruffaerts, R., … & Kessler, R. C. (2015). Psychotic experiences in the general population: a cross-national analysis based on 31 261 respondents from 18 countries. JAMA psychiatry, 72(7), 697–705.

12. Van Os, J., Linscott, R. J., Myin-Germeys, I., Delespaul, P., & Krabbendam, L. J. P. M. (2009). A systematic review and meta-analysis of the psychosis continuum: evidence for a psychosis proneness-persistence-impairment model of psychotic disorder. Psychological medicine, 39(2), 179.

13. Baumeister, D., Sedgwick, O., Howes, O., & Peters, E. (2017). Auditory verbal hallucinations and continuum models of psychosis: a systematic review of the healthy voice-hearer literature. Clinical psychology review, 51, 125–141.

14. Northoff, G. (2014). Are auditory hallucinations related to the brain’s resting state activity? A neurophenomenal resting state hypothesis. Clinical Psychopharmacology and Neuroscience, 12(3), 189.

15. Alderson-Day, B., McCarthy-Jones, S., & Fernyhough, C. (2015). Hearing voices in the resting brain: a review of intrinsic functional connectivity research on auditory verbal hallucinations. Neuroscience & Biobehavioral Reviews, 55, 78–87.

16. Alderson-Day, B., Diederen, K., Fernyhough, C., Ford, J. M., Horga, G., Margulies, D. S., … & Jardri, R. (2016). Auditory hallucinations and the brain’s resting-state networks: findings and methodological observations. Schizophrenia bulletin, 42(5), 1110–1123.

17. Ćurčić-Blake, B., Ford, J. M., Hubl, D., Orlov, N. D., Sommer, I. E., Waters, F., … & Aleman, A. (2017). Interaction of language, auditory and memory brain networks in auditory verbal hallucinations. Progress in neurobiology, 148, 1–20.

18. Verdoux, H., & van Os, J. (2002). Psychotic symptoms in non-clinical populations and the continuum of psychosis. Schizophrenia research, 54(1-2), 59–65.

19. Kelleher, I., Jenner, J. A., & Cannon, M. (2010). Psychotic symptoms in the general population–an evolutionary perspective. The British Journal of Psychiatry, 197(3), 167–169.

20. Kelleher, I., & Cannon, M. (2011). Psychotic-like experiences in the general population: characterizing a high-risk group for psychosis. Psychological medicine, 41(1), 1–6.

21. Leng, B., Han, S., Bao, Y., Zhang, H., Wang, Y., Wu, Y., & Wang, Y. (2016). The uncinate fasciculus as observed using diffusion spectrum imaging in the human brain. Neuroradiology, 58(6), 595–606.

22. Hau, J., Sarubbo, S., Houde, J. C., Corsini, F., Girard, G., Deledalle, C., … & Petit, L. (2017). Revisiting the human uncinate fasciculus, its subcomponents and asymmetries with stem-based tractography and microdissection validation. Brain Structure and Function, 222(4), 1645–1662.

23. Briggs, R. G., Rahimi, M., Conner, A. K., Sali, G., Baker, C. M., Burks, J. D., … & Sughrue, M. E. (2018). A Connectomic Atlas of the Human Cerebrum—Chapter 15: Tractographic Description of the Uncinate Fasciculus. Operative Neurosurgery, 15(suppl_1), S450-S455.

24. Schmahmann, J. D., Smith, E. E., Eichler, F. S., & Filley, C. M. (2008). Cerebral white matter: neuroanatomy, clinical neurology, and neurobehavioral correlates. Annals of the New York Academy of Sciences, 1142, 266.

25. Von Der Heide, R. J., Skipper, L. M., Klobusicky, E., & Olson, I. R. (2013). Dissecting the uncinate fasciculus: disorders, controversies, and a hypothesis. Brain, 136(6), 1692–1707.

26. Iwabuchi, S. J., Liddle, P. F., & Palaniyappan, L. (2015). Structural connectivity of the salience-executive loop in schizophrenia. European archives of psychiatry and clinical neuroscience, 265(2), 163–166.

27. Olson, I. R., Von Der Heide, R. J., Alm, K. H., & Vyas, G. (2015). Development of the uncinate fasciculus: Implications for theory and developmental disorders. Developmental cognitive neuroscience, 14, 50–61.

28. Szeszko, P. R., Robinson, D. G., Ashtari, M., Vogel, J., Betensky, J., Sevy, S., … & Bilder, R. M. (2008). Clinical and neuropsychological correlates of white matter abnormalities in recent onset schizophrenia. Neuropsychopharmacology, 33(5), 976–984.

29. Kitis, O., Ozalay, O., Zengin, E. B., Haznedaroglu, D., Eker, M. C., Yalvac, D., … & Gonul, A. S. (2012). Reduced left uncinate fasciculus fractional anisotropy in deficit schizophrenia but not in non-deficit schizophrenia. Psychiatry and clinical neurosciences, 66(1), 34–43.

30. Voineskos, A. N., Foussias, G., Lerch, J., Felsky, D., Remington, G., Rajji, T. K., … & Mulsant, B. H. (2013). Neuroimaging evidence for the deficit subtype of schizophrenia. JAMA psychiatry, 70(5), 472–480.

31. Lei, W., Li, N., Deng, W., Li, M., Huang, C., Ma, X., … & Li, T. (2015). White matter alterations in first episode treatment-naive patients with deficit schizophrenia: a combined VBM and DTI study. Scientific reports, 5(1), 1–11.

32. Mandl, R. C., Schnack, H. G., Luigjes, J., Van Den Heuvel, M. P., Cahn, W., Kahn, R. S., & Hulshoff Pol, H. E. (2010). Tract-based analysis of magnetization transfer ratio and diffusion tensor imaging of the frontal and frontotemporal connections in schizophrenia. Schizophrenia bulletin, 36(4), 778–787.

33. Sun, H., Lui, S., Yao, L., Deng, W., Xiao, Y., Zhang, W., … & Gong, Q. (2015). Two patterns of white matter abnormalities in medication-naive patients with first-episode schizophrenia revealed by diffusion tensor imaging and cluster analysis. JAMA psychiatry, 72(7), 678–686.

34. Luck, D., Buchy, L., Czechowska, Y., Bodnar, M., Pike, G. B., Campbell, J. S., … & Lepage, M. (2011). Fronto-temporal disconnectivity and clinical short-term outcome in first episode psychosis: a DTI-tractography study. Journal of psychiatric research, 45(3), 369–377.

35. Hubl, D., Koenig, T., Strik, W., Federspiel, A., Kreis, R., Boesch, C., … & Dierks, T. (2004). Pathways that make voices: white matter changes in auditory hallucinations. Archives of general psychiatry, 61(7), 658–668.

36. De Weijer, A. D., Mandl, R. C. W., Diederen, K. M. J., Neggers, S. F. W., Kahn, R. S., Pol, H. H., & Sommer, I. E. C. (2011). Microstructural alterations of the arcuate fasciculus in schizophrenia patients with frequent auditory verbal hallucinations. Schizophrenia research, 130(1-3), 68–77.

37. De Weijer, A. D., Neggers, S. F., Diederen, K. M., Mandl, R. C., Kahn, R. S., Hulshoff Pol, H. E., & Sommer, I. E. (2013). Aberrations in the arcuate fasciculus are associated with auditory verbal hallucinations in psychotic and in non-psychotic individuals. Human brain mapping, 34(3), 626–634.

38. Curčić-Blake, B., Nanetti, L., van der Meer, L., Cerliani, L., Renken, R., Pijnenborg, G. H., & Aleman, A. (2015). Not on speaking terms: hallucinations and structural network disconnectivity in schizophrenia. Brain Structure and Function, 220(1), 407–418.

39. Filippi, M., Canu, E., Gasparotti, R., Agosta, F., Valsecchi, P., Lodoli, G., … & Sacchetti, E. (2014). Patterns of brain structural changes in first-contact, antipsychotic drug-naive patients with schizophrenia. American Journal of Neuroradiology, 35(1), 30–37.

40. Bopp, M. H., Zöllner, R., Jansen, A., Dietsche, B., Krug, A., & Kircher, T. T. (2017). White matter integrity and symptom dimensions of schizophrenia: a diffusion tensor imaging study. Schizophrenia research, 184, 59–68.

41. Jung, S., Kim, J. H., Sung, G., Ko, Y. G., Bang, M., Park, C. I., & Lee, S. H. (2020). Uncinate fasciculus white matter connectivity related to impaired social perception and cross-sectional and longitudinal symptoms in patients with schizophrenia spectrum psychosis. Neuroscience Letters, 737, 135144.

42. Ćurčić-Blake, B., Houenou, J., & Jardri, R. (2018). Dysconnectivity in Hallucinations. In Hallucinations in Psychoses and Affective Disorders (pp. 159-171). Springer, Cham.

43. Frith, C. D., & Done, D. J. (1988). Towards a neuropsychology of schizophrenia. The British Journal of Psychiatry, 153(4), 437–443.

44. Allen, P., Aleman, A., & Mcguire, P. K. (2007). Inner speech models of auditory verbal hallucinations: evidence from behavioural and neuroimaging studies. International Review of Psychiatry, 19(4), 407–415.

45. Whitford, T. J., Ford, J. M., Mathalon, D. H., Kubicki, M., & Shenton, M. E. (2012). Schizophrenia, myelination, and delayed corollary discharges: a hypothesis. Schizophrenia bulletin, 38(3), 486–494.

46. Catani, M., Craig, M. C., Forkel, S. J., Kanaan, R., Picchioni, M., Toulopoulou, T., … & McGuire, P. (2011). Altered integrity of perisylvian language pathways in schizophrenia: relationship to auditory hallucinations. Biological psychiatry, 70(12), 1143–1150.

47. McCarthy-Jones, S., Oestreich, L. K., Bank, A. S. R., & Whitford, T. J. (2015). Reduced integrity of the left arcuate fasciculus is specifically associated with auditory verbal hallucinations in schizophrenia. Schizophrenia research, 162(1-3), 1–6.

48. Psomiades, M., Fonteneau, C., Mondino, M., Luck, D., Haesebaert, F., Suaud-Chagny, M. F., & Brunelin, J. (2016). Integrity of the arcuate fasciculus in patients with schizophrenia with auditory verbal hallucinations: A DTI-tractography study. NeuroImage: Clinical, 12, 970–975.

49. Leroux, E., Delcroix, N., & Dollfus, S. (2017). Abnormalities of language pathways in schizophrenia patients with and without a lifetime history of auditory verbal hallucinations: A DTI-based tractography study. The World Journal of Biological Psychiatry, 18(7), 528–538.

50. Chawla, N., Deep, R., Khandelwal, S. K., & Garg, A. (2019). Reduced integrity of superior longitudinal fasciculus and arcuate fasciculus as a marker for auditory hallucinations in schizophrenia: a DTI tractography study. Asian journal of psychiatry, 44, 179–186.

51. Rotarska-Jagiela, A., Oertel-Knoechel, V., DeMartino, F., van de Ven, V., Formisano, E., Roebroeck, A., … & Linden, D. E. (2009). Anatomical brain connectivity and positive symptoms of schizophrenia: a diffusion tensor imaging study. Psychiatry Research: Neuroimaging, 174(1), 9–16.

52. Catani, M., & De Schotten, M. T. (2008). A diffusion tensor imaging tractography atlas for virtual in vivo dissections. Cortex, 44(8), 1105–1132.

53. Eddy, C. M. (2016). The junction between self and other? Temporo-parietal dysfunction in neuropsychiatry. Neuropsychologia, 89, 465–477.

54. Quesque, F., & Brass, M. (2019). The role of the temporoparietal junction in self-other distinction. Brain topography, 32(6), 943–955.

55. Steinmann, S., Leicht, G., & Mulert, C. (2014). Interhemispheric auditory connectivity: structure and function related to auditory verbal hallucinations. Frontiers in human neuroscience, 8, 55.

56. Steinmann, S., Leicht, G., & Mulert, C. (2019). The interhemispheric miscommunication theory of auditory verbal hallucinations in schizophrenia. International Journal of Psychophysiology, 145, 83–90.

57. Shergill, S. S., Kanaan, R. A., Chitnis, X. A., O’Daly, O., Jones, D. K., Frangou, S., … & McGuire, P. (2007). A diffusion tensor imaging study of fasciculi in schizophrenia. American Journal of Psychiatry, 164(3), 467–473.

58. Mulert, C., Kirsch, V., Whitford, T. J., Alvarado, J., Pelavin, P., McCarley, R. W., … & Shenton, M. E. (2012). Hearing voices: a role of interhemispheric auditory connectivity? The World Journal of Biological Psychiatry, 13(2), 153–158.

59. Wigand, M., Kubicki, M., Clemm von Hohenberg, C., Leicht, G., Karch, S., Eckbo, R., … & Mulert, C. (2015). Auditory verbal hallucinations and the interhemispheric auditory pathway in chronic schizophrenia. The World Journal of Biological Psychiatry, 16(1), 31–44.

60. Xi, Y. B., Guo, F., Li, H., Chang, X., Sun, J. B., Zhu, Y. Q., … & Yin, H. (2016). The structural connectivity pathology of first-episode schizophrenia based on the cardinal symptom of auditory verbal hallucinations. Psychiatry Research: Neuroimaging, 257, 25–30.

61. Zhang, X., Gao, J., Zhu, F., Wang, W., Fan, Y., Ma, Q., … & Yang, J. (2018). Reduced white matter connectivity associated with auditory verbal hallucinations in first episode and chronic schizophrenia: a diffusion tensor imaging study. Psychiatry Research: Neuroimaging, 273, 63–70.

62. Whitford, T. J., Kubicki, M., Schneiderman, J. S., O’Donnell, L. J., King, R., Alvarado, J. L., … & Shenton, M. E. (2010). Corpus callosum abnormalities and their association with psychotic symptoms in patients with schizophrenia. Biological psychiatry, 68(1), 70–77.

63. Hofer, S., & Frahm, J. (2006). Topography of the human corpus callosum revisited— comprehensive fiber tractography using diffusion tensor magnetic resonance imaging. Neuroimage, 32(3), 989–994.

64. Jordan, M. I., & Rumelhart, D. E. (1992). Forward models: Supervised learning with a distal teacher. Cognitive science, 16(3), 307–354.

65. Friston, K. (2012). Prediction, perception, and agency. International Journal of Psychophysiology, 83(2), 248–252.

66. Wolpert, D. M., Miall, R. C., & Kawato, M. (1998). Internal models in the cerebellum. Trends in cognitive sciences, 2(9), 338–347.

67. Tanaka, H., Ishikawa, T., Lee, J., & Kakei, S. (2020). The cerebro-cerebellum as a locus of forward model: a review. Frontiers in systems neuroscience, 14, 19.

68. Pinheiro, A. P., Schwartze, M., & Kotz, S. A. (2020). Cerebellar circuitry and auditory verbal hallucinations: An integrative synthesis and perspective. Neuroscience & Biobehavioral Reviews.

69. Larøi, F., & Van Der Linden, M. (2005). Nonclinical Participants’ Reports of Hallucinatory Experiences. Canadian Journal of Behavioural Science/Revue canadienne des sciences du comportement, 37(1), 33.

70. Smith, S. M., Jenkinson, M., Woolrich, M. W., Beckmann, C. F., Behrens, T. E., Johansen-Berg, H., … & Matthews, P. M. (2004). Advances in functional and structural MR image analysis and implementation as FSL. Neuroimage, 23, S208–S219.

71. Andersson, J. L., Skare, S., & Ashburner, J. (2003). How to correct susceptibility distortions in spin-echo echo-planar images: application to diffusion tensor imaging. Neuroimage, 20(2), 870–888.

72. Smith, S. M. (2002). Fast robust automated brain extraction. Human brain mapping, 17(3), 143–155.

73. Andersson, J. L., & Sotiropoulos, S. N. (2016). An integrated approach to correction for off-resonance effects and subject movement in diffusion MR imaging. Neuroimage, 125, 1063–1078.

74. Behrens, T. E., Woolrich, M. W., Jenkinson, M., Johansen-Berg, H., Nunes, R. G., Clare, S., … & Smith, S. M. (2003). Characterization and propagation of uncertainty in diffusion-weighted MR imaging. Magnetic Resonance in Medicine: An Official Journal of the International Society for Magnetic Resonance in Medicine, 50(5), 1077–1088.

75. Jenkinson, M., Bannister, P., Brady, M., & Smith, S. (2002). Improved optimization for the robust and accurate linear registration and motion correction of brain images. Neuroimage, 17(2), 825–841.

76. Hernandez-Fernandez, M., Reguly, I., Jbabdi, S., Giles, M., Smith, S., & Sotiropoulos, S. N. (2019). Using GPUs to accelerate computational diffusion MRI: From microstructure estimation to tractography and connectomes. Neuroimage, 188, 598–615.

77. Alexander, A. L., Lee, J. E., Lazar, M., & Field, A. S. (2007). Diffusion tensor imaging of the brain. Neurotherapeutics, 4(3), 316–329.

78. Smith, S. M., Jenkinson, M., Johansen-Berg, H., Rueckert, D., Nichols, T. E., Mackay, C. E., … & Behrens, T. E. (2006). Tract-based spatial statistics: voxelwise analysis of multi-subject diffusion data. Neuroimage, 31(4), 1487–1505.

79. Hau, J., Sarubbo, S., Perchey, G., Crivello, F., Zago, L., Mellet, E., … & Petit, L. (2016). Cortical terminations of the inferior fronto-occipital and uncinate fasciculi: anatomical stem-based virtual dissection. Frontiers in neuroanatomy, 10, 58.

80. Hau, J., Sarubbo, S., Houde, J. C., Corsini, F., Girard, G., Deledalle, C., … & Petit, L. (2017). Revisiting the human uncinate fasciculus, its subcomponents and asymmetries with stem-based tractography and microdissection validation. Brain Structure and Function, 222(4), 1645–1662.

81. O’Hanlon, E., Leemans, A., Kelleher, I., Clarke, M. C., Roddy, S., Coughlan, H., … & Cannon, M. (2015). White matter differences among adolescents reporting psychotic experiences: a population-based diffusion magnetic resonance imaging study. JAMA psychiatry, 72(7), 668–677.

82. Boos, H. B., Mandl, R. C., van Haren, N. E., Cahn, W., van Baal, G. C. M., Kahn, R. S., & Pol, H. E. H. (2013). Tract-based diffusion tensor imaging in patients with schizophrenia and their non-psychotic siblings. European Neuropsychopharmacology, 23(4), 295–304.

83. Menon, V. (2011). Large-scale brain networks and psychopathology: a unifying triple network model. Trends in cognitive sciences, 15(10), 483–506.

84. Palaniyappan, L., & Liddle, P. F. (2012). Does the salience network play a cardinal role in psychosis? An emerging hypothesis of insular dysfunction. Journal of Psychiatry & Neuroscience.

85. Northoff G. (2014). Are Auditory Hallucinations Related to the Brain’s Resting State Activity? A ‘Neurophenomenal Resting State Hypothesis’. Clinical psychopharmacology and neuroscience: the official scientific journal of the Korean College of Neuropsychopharmacology, 12(3), 189–195.

86. Robinson, J. D., Wagner, N. F., & Northoff, G. (2016). Is the sense of agency in schizophrenia influenced by resting-state variation in self-referential regions of the brain? Schizophrenia bulletin, 42(2), 270–276.

87. Wotruba, D., Michels, L., Buechler, R., Metzler, S., Theodoridou, A., Gerstenberg, M., … & Heekeren, K. (2014). Aberrant coupling within and across the default mode, task-positive, and salience network in subjects at risk for psychosis. Schizophrenia bulletin, 40(5), 1095–1104.

88. Bolton, T. A., Wotruba, D., Buechler, R., Theodoridou, A., Michels, L., Kollias, S., … & Van De Ville, D. (2020). Triple network model dynamically revisited: lower salience network state switching in pre-psychosis. Frontiers in physiology, 11, 66.

89. Seok, J. H., Park, H. J., Chun, J. W., Lee, S. K., Cho, H. S., Kwon, J. S., & Kim, J. J. (2007). White matter abnormalities associated with auditory hallucinations in schizophrenia: a combined study of voxel-based analyses of diffusion tensor imaging and structural magnetic resonance imaging. Psychiatry Research: Neuroimaging, 156(2), 93–104.

90. Friston, K. J. (2005). Hallucinations and perceptual inference. Behav Brain Sci, 28(6), 764–766.

91. Hugdahl, K. (2009). “Hearing voices”: Auditory hallucinations as failure of top-down control of bottom-up perceptual processes. Scandinavian journal of psychology, 50(6), 553–560.

92. Hugdahl, K. (2017). Auditory hallucinations as translational psychiatry: Evidence from magnetic resonance imaging. Balkan medical journal, 34(6), 504.

93. Sterzer, P., Adams, R. A., Fletcher, P., Frith, C., Lawrie, S. M., Muckli, L., … & Corlett, P. R. (2018). The predictive coding account of psychosis. Biological psychiatry, 84(9), 634–643.

94. Johnson, J. F., Belyk, M., Schwartze, M., Pinheiro, A. P., & Kotz, S. A. (2019). The role of the cerebellum in adaptation: ALE meta-analyses on sensory feedback error. Human brain mapping, 40(13), 3966–3981.

95. Johnson JF, Belyk M, Schwartze M, Pinheiro AP, Kotz SA (2021). Expectancy changes the self-monitoring of voice identity. European Journal of Neuroscience, 53(8), 2681–2695.

96. Leroux, E., Delcroix, N., & Dollfus, S. (2015). Left-hemisphere lateralization for language and interhemispheric fiber tracking in patients with schizophrenia. Schizophrenia research, 165(1), 30–37.

97. Miall, R. C., Weir, D. J., Wolpert, D. M., & Stein, J. F. (1993). Is the cerebellum a smith predictor?. Journal of motor behavior, 25(3), 203–216.

98. Welniarz, Q., Worbe, Y., & Gallea, C. (2021). The forward model: a unifying theory for the role of the cerebellum in motor control and sense of agency. Frontiers in Systems Neuroscience, 15.

99. Kim, D. J., Kent, J. S., Bolbecker, A. R., Sporns, O., Cheng, H., Newman, S. D., … & Hetrick, W. P. (2014). Disrupted modular architecture of cerebellum in schizophrenia: a graph theoretic analysis. Schizophrenia bulletin, 40(6), 1216–1226.

100. Zhang, X. Y., Fan, F. M., Chen, D. C., Tan, Y. L., Tan, S. P., Hu, K., … & Soares, J. C. (2016). Extensive white matter abnormalities and clinical symptoms in drug-naive patients with first-episode schizophrenia: a voxel-based diffusion tensor imaging study. The Journal of clinical psychiatry, 77(2), 205–211.

