## Supplementary Information for "Variability in white matter structure relates to hallucination proneness"

| **Supplementary Table 1. Atlas Information** | | | |
| --- | --- | --- | --- |
| ***A. Atlas-Based ROI Masks*** | | | |
| Pathway | | | Atlas |
| L UF | | | JHU-white matter tractography |
| R UF | | | JHU-white matter tractography |
| L AF A | | | NATRAINLAB |
| R AF A | | | NATRAINLAB |
| L AF L | | | NATRAINLAB |
| R AF L | | | NATRAINLAB |
| L AF P | | | NATRAINLAB |
| R AF P | | | NATRAINLAB |
| CC B | | | JHU ICBM-DTI-81 white matter |
| CC G | | | JHU ICBM-DTI-81 white matter |
| CC S | | | JHU ICBM-DTI-81 white matter |
| ***B. Tractography Guiding Masks*** | | | |
| Pathway | Type | Region | Atlas |
| CTC | Seed | CE (R/L) | FSL Cerebellar Atlas in MNI152 |
|  | Inclusion | sCEP (IL) | JHU ICBM-DTI-81 white matter |
|  |  | VLTN (CL) | Talaraich Daeomon |
|  | Exclusion | mCEP | JHU ICBM-DTI-81 white matter |
| CPC | Seed | M1 (R/L) | Juelich Histological |
|  | Inclusion | CP* (IL) | JHU ICBM-DTI-81 white matter |
|  |  | Pons | Talaraich Daeomon |
|  |  | mCEP (CL) | JHU ICBM-DTI-81 white matter |
|  |  | CE (CL) | FSL Cerebellar Atlas in MNI152 |
|  | Exclusion | CE (IL) | FSL Cerebellar Atlas in MNI152 |
| Legend: A. Atlas-based region of interest masks: R = right, L = left, UF = uncinate fasciculus, AF = arcuate fasciculus (A = anterior, L = longitudinal, P = posterior), CC = corpus callosum (B = body, G = genus, S = splenium); B. Tractography guiding masks: CE = cerebellum, CEP = cerebellar peduncle (s = superior, m = middle), VLTN = ventrolateral thalamic nuclei, M1 = primary motor cortex, CP = cerebral peduncle (* = posterior corticopontine segment of the crus cerebri) Pons = pontine nuclei, Midline = hemispheric crossing before decussation point, IL = ipsilateral, CL = contralateral | | | |

| **Supplementary Table 2. LSHS Auditory Item-Subscale Correlation Results** | | | | |
| --- | --- | --- | --- | --- |
| Tract | | *r* | 95% CI (bias-corrected) | FDR-*p* |
| ***A. Atlas-based ROIs: FA correlation*** | | | | |
| Arcuate Fasciculus | | | | |
| Left | Longitudinal | 0.120 | -0.129-0.415 | 0.457 |
|  | Anterior | 0.153 | -0.066-0.422 | 0.387 |
|  | Posterior | 0.197 | -0.041-0.409 | 0.267 |
| Right | Longitudinal | 0.047 | -0.188-0.303 | 0.745 |
|  | Anterior | 0.202 | -0.050-0.486 | 0.267 |
|  | Posterior | 0.259 | 0.000-0.550 | 0.158 |
| Uncinate Fasciculus | | | | |
| Left | | 0.145 | -0.120-0.433 | 0.387 |
| Right | | 0.279 | 0.026-0.532 | 0.158 |
| Corpus Callosum | | | | |
| Genus | | 0.260 | -0.010-0.497 | 0.158 |
| Body | | 0.350 | 0.099-0.597 | 0.143 |
| Splenium | | 0.257 | -0.005-0.495 | 0.072 |
| ***B. Tractography-based ROIs: FA and Streamline correlation*** | | | | |
| Cortico-ponto-cerebellar Pathway | | | |  |
| FA | Right → Left | 0.095 | -0.206-0.359 | 0.793 |
|  | Left → Right | 0.134 | -0.163-0.393 | 0.793 |
| # SL | Right → Left | 0.055 | -0.234-0.379 | 0.718 |
|  | Left → Left | -0.100 | -0.312-0.144 | 0.676 |
| Cerebello-thalamo-cortical Pathway | | | |  |
| FA | Left → Right | -0.083 | -0.322-0.153 | 0.793 |
|  | Right → Left | 0.043 | -0.341-0.340 | 0.793 |
| # SL | Left → Right | 0.308 | -0.051-0.618 | 0.112 |
|  | Right → Left | 0.369 |  | 0.076 |
| Legend: Pearson’s correlation analysis with hallucination proneness. Benjamini-Hochberg FDR-p correction for multiple comparisons (* = p < 0.05 threshold for significance). Bias-corrected confidence intervals (95%, bootstrapping with 1000 samples). | | | | |
